# Unique unbiased median solution for even sample sizes

**DOI:** 10.1101/2025.08.05.668614

**Authors:** Mikhail Y. Lipin, Elias Benjamin Crampton, Steven A. Thomas

## Abstract

Data in experimental biology are frequently marred by outliers and asymmetric distributions. The median, being a robust estimator of central tendency, is less sensitive to outliers than the mean. However, for ranked datasets with an even number of observations, the conventional median—calculated as the average of the two middle values—can introduce bias by implicitly assuming symmetry in the data distribution. This study aims to identify a median estimator that is unbiased.

To derive the unbiased median estimator, we minimized the sum of residuals raised to a rational power approaching one. We compared the properties of the unbiased and conventional medians using Poisson-distributed datasets. Random samples were generated with the Mersenne Twister algorithm implemented in IgorPro software (WaveMetrics Inc., Oregon).

For odd sample sizes, the unbiased median coincides with the conventional median (the middle value). For even sample sizes, the unbiased median is defined as the value that equalizes the product of distances to data points above and below it—a definition that differs from the conventional median in asymmetric distributions. Although both median estimators tend to underestimate the mean of Poisson-distributed data, the unbiased median is consistently closer to the expected value. Additionally, the unbiased median exhibits lower variance compared to the conventional median.

Thus, for even sample sizes, the proposed unbiased median provides a central tendency measure that is unbiased, more accurate, and has reduced variance relative to the conventional median.

## Introduction

The concept of the median was first introduced in 1755 by the distinguished Croatian physicist, astronomer, and mathematician Roger Joseph Boscovich as a method for fitting a line to astronomical observations [1]. Boscovich sought the value that minimizes the sum of absolute deviations, discovering that this value separates the lower half of a dataset from the upper half. For ordinal data—where elements are ranked relative to one another but not necessarily measured on an absolute scale—only the median and mode serve as appropriate measures of central tendency [2]. Consequently, the median is an especially valuable statistical measure for biological data, which are often ordinal, asymmetrically distributed, and contain outliers.

The median proves to be a powerful tool in biological data analysis whenever a robust measure of central tendency is required to represent a “typical” value, such as cell size, metabolite concentration, or dispersal distance [3]. It is also commonly used to determine typical survival times in populations [4]. In pharmacology, median effective dose (ED50) and median toxic dose (TD50)—which indicate doses causing specific effects or toxicity in 50% of a population—are essential for assessing drug safety and efficacy [5].

For datasets with an odd number of samples, the median—defined as the value minimizing the sum of absolute deviations—is simply the middle value. However, with an even number of samples, this approach leads to an ill-posed problem: the solution is not unique, as any point between the two middle values minimizes the sum of absolute deviations. Notably, there is no historical record of when or who first defined the median for even-sized samples as the average of the two middle values. Moreover, this conventional definition is poorly justified and only appropriate under the strong assumption of a symmetric data distribution—an assumption rarely valid for most biological datasets.

In this study, we demonstrate that the problem of defining the median for even sample sizes can be well-posed; specifically, the location of the median can be uniquely determined without bias toward symmetrical distributions. To assess the performance of this unbiased median, we compared it with the conventional median by measuring the central tendency of Poisson-distributed datasets. The Poisson process was chosen for its fundamental role in modeling random independent events in biology, such as genetic mutations [6], cell divisions [7], and cancer development [8]. Our results show that the unbiased median provides a more accurate estimate of central tendency and exhibits lower variance than the conventional median.

## Methods and Results

Jacques Hadamard defined a problem as *well-posed* if its solution exists, is unique, and depends continuously on the input data [9]. These three criteria have since become fundamental concepts in mathematics. In contrast, finding the median for an even number of samples by minimizing the sum of absolute deviations is an *ill-posed* problem because the solution is not unique and does not depend continuously on the data. However, by framing this ill-posed problem as a limiting case of a well-posed problem—specifically, by minimizing the sum of residuals raised to the power of a rational number that approaches one—we can obtain a solution that is both unique and depends continuously on the data.

Formally, consider a ranked dataset *X* = {*x*_1_ < *x*_2_ < … *x*_*M*_} consisting of *M* measurements.

We seek an estimate *α* that minimizes the objective function:

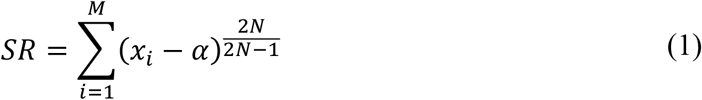

where *N* is a natural number.

The even numerator 2*N* in the exponent ensures that each term in the sum (1) is non-negative. To find the minimum of the sum of residuals *SR* defined by equation (1), we take its first derivative with respect to *α*:

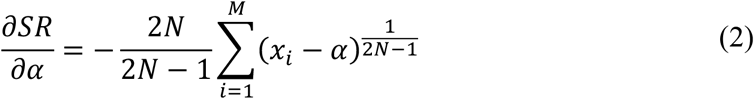

and set it equal to zero:

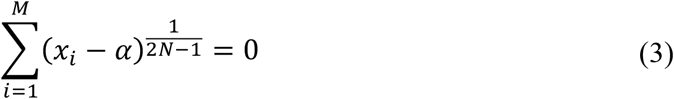

In a special case where *N* = 1, the equation (3) holds true when the estimate *α* equals the arithmetic mean of the dataset:

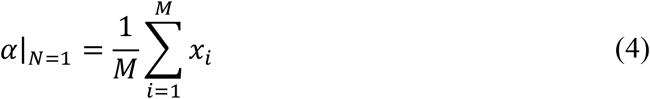

The mean provides the best estimate when the distribution of the dataset *X* is symmetrical. For asymmetrical distributions, we transform each term in the derivative (equation 3) as follows:

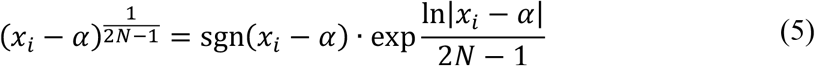

where the sign function “sgn(_*_)” is:

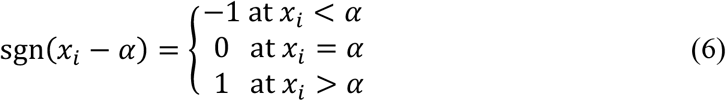

For large values of *N* (*N* ≫ 1), the exponent in equation (5) can be expanded into a Taylor series:

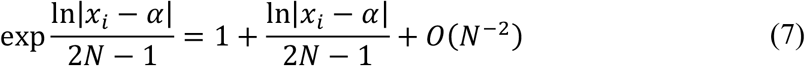

Therefore, each term in the sum from equation (3) can be approximated as:

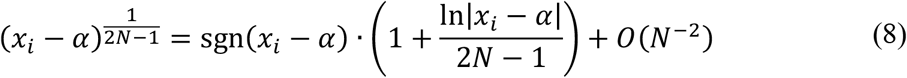

Neglecting terms of order *O*(*N*^−2^) for *N* ≫ 1, and substituting these approximations back into equation (3), the condition reduces to:

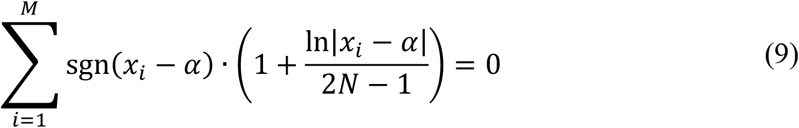

For an even number of samples *M* = 2*K*, the equation (9) holds true at *x* _*K*_ < *α* < *x* _*K*+1_:

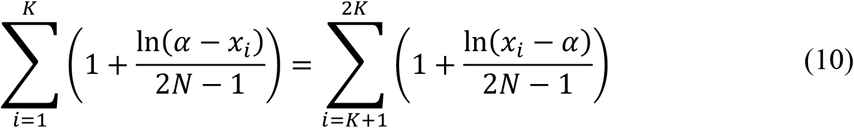

As the numbers of ones in the sums of left and right sides of the equation (10) are equal, they cancel each other, and the equation simplifies to:

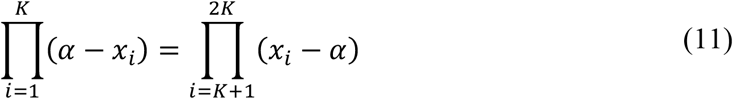

from which the estimate *α* at *K* = 1, 2 can be found as follows:

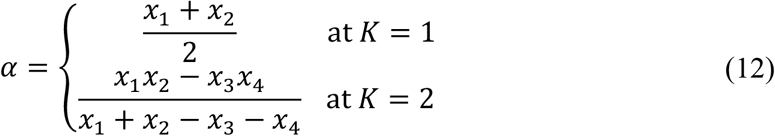

At *K* = 3 to 4, the estimate *α* is a root of the third-degree polynomial, whereas at *K* ≥ 5 *α* can be calculated iteratively.

For an odd number of samples *M* = 2*K*+1, the equation (9) can be rearranged into:

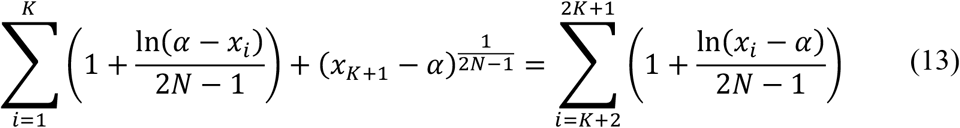

from which the residual *x*_*K*+1_ – *α* is found as:

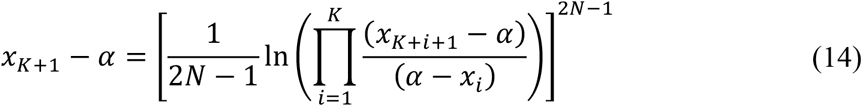

At *N* ≫ 1, the right part of the equation (14) approaches zero, thus leaving:

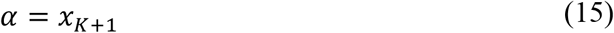

that matches the conventional median of the dataset with an odd number of samples.

Note that at *N* ⟶ ∞ the equation (1) can be simplified:

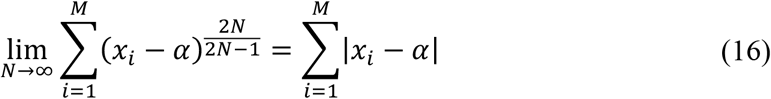

Thus, as *N* ⟶ ∞, the estimate *α* minimizes the sum of distances to the values in dataset *X*. Since this estimate *α* from equation (11) is not biased toward the arithmetic mean of the middle values and minimizes the sum of distances (equation 16), we refer to it as the “unbiased median”.

To evaluate the performance of the proposed unbiased median on datasets with asymmetric distributions, we performed computer simulations of Poisson processes with probability mass function *f*(*k*; λ) :

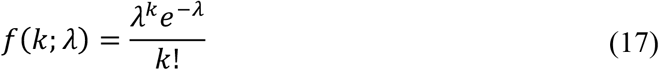

where *k* denotes the number of events and λ is the expected value. Random Poisson-distributed samples were generated using IgorPro software (WaveMetrics Inc., Oregon), which employs the Mersenne Twister pseudo-random number generator. The unbiased median *α* was calculated in IgorPro by numerically solving equation (11) with an accuracy of 10^−6^.

To illustrate the differences among the unbiased median, conventional median, and mean, we plotted their corresponding central tendency estimates for datasets with varying sample sizes (Fig 1). Despite the considerable noise inherent to the Poisson distribution, it is evident that the unbiased median 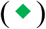 often either coincides with the conventional median 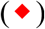—notably at sample sizes n = 10, 12, 14 in panel ***A*** and n = 8, 10 in panel ***B***—or tends to shift closer to the mean (♦). Importantly, the unbiased median frequently aligns more closely with the mean.

**Fig 1.**
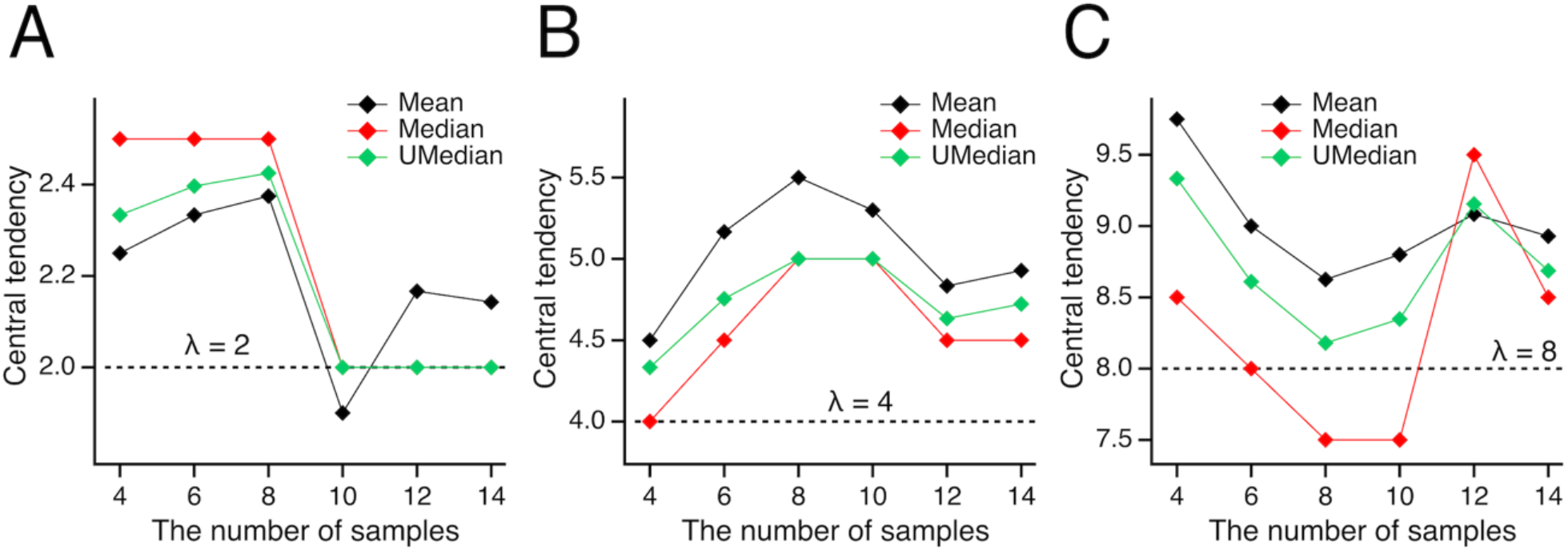
Unbiased median vs. conventional median and mean. Central tendency estimates for Poisson-distributed datasets—evaluated as mean (♦), unbiased median 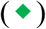, and conventional median 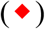—are plotted against sample size for expected values λ of 2, 4, and 8 (dashed lines) in panels ***A, B***, and ***C***, respectively. Notably, the unbiased median either coincides with the conventional median (e.g., at sample sizes *n* = 10, 12, 14 in panel ***A*** and *n* = 8 and 10 in panel ***B***) or is shifted toward the mean, highlighting its tendency to align more closely with the mean value as sample characteristics vary.

The tendency of the unbiased median to lean toward the mean suggests that its values may be closer to the mean. To evaluate this hypothesis, we conducted a large number of simulations (n = 3·10^7^) and computed the arithmetic means of the central tendency estimates to reduce noise. Here and in subsequent analyses, we consider the arithmetic means of three estimators: the arithmetic mean (♦), the conventional median 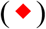, and the unbiased median 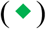 (Fig 2). As anticipated for the Poisson process, the arithmetic mean (♦) closely matches the expected value λ (Fig 2), with standard deviations (SD) from λ of 1.9·10^−5^, 2.7·10^−5^, 4.1·10^−5^, 3.6·10^−5^, and 5.0·10^−5^ for λ = 1, 2, 3, 4, and 5, respectively. Since all these SDs are greater than 10^−6^, which is the numerical precision for calculating the unbiased median, they set the lower bound for inaccuracy in subsequent analyses. Importantly, compared to the conventional median 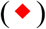, the unbiased median 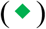 consistently provides estimates that are nearer to the expected value λ.

**Fig 2.**
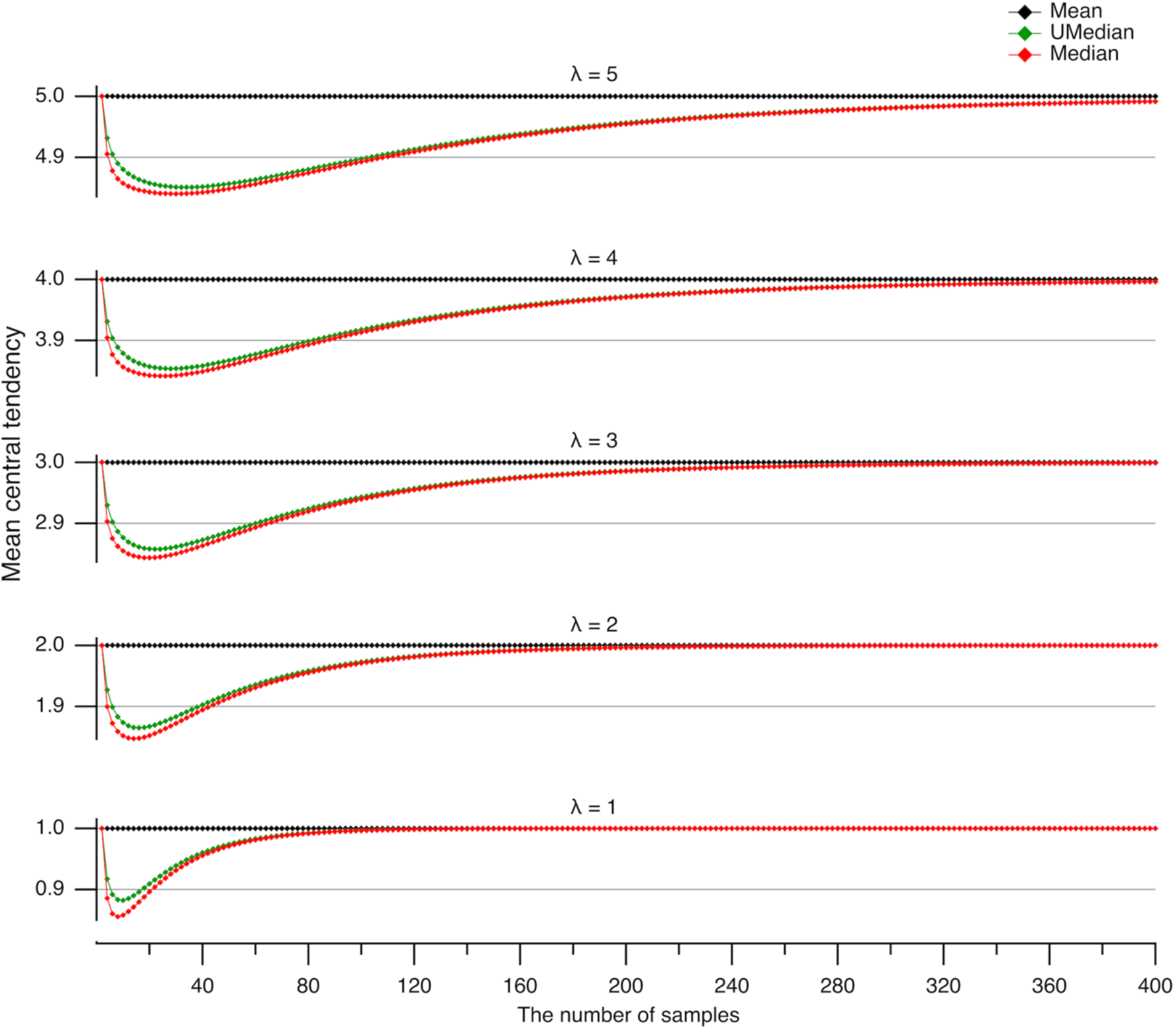
Underestimation of the expected value by medians. The arithmetic mean (calculated from 3·10^7^ simulations) of central tendency estimates for Poisson-distributed datasets with expected values λ ranging from 1 to 5 (bottom to top) is plotted against the number of samples. Central tendencies are evaluated as the mean (♦), unbiased median 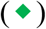, and conventional median 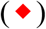. Note that both median estimators tend to underestimate the expected value λ, but the unbiased median consistently provides estimates closer to the mean than the conventional median.

Notably, the central tendency measured by the medians is significantly underestimated for small sample sizes (Fig 2) because medians are insensitive to the extreme positive values that are intrinsic to the Poisson process. These extreme values are not outliers; rather, they are a natural part of the distribution. As a result, the medians’ disregard of these values leads to an underestimation of the true central tendency. Therefore, using the mean—which correctly estimates the expected value of a Poisson-distributed dataset—is more appropriate here than using the median, which tends to underestimate it.

However, if these extreme values were true outliers, the median would provide a more accurate measure of central tendency than the mean. Thus, the underestimation of the mean by the medians (see Fig 2) reflects their ability to mitigate the influence of extreme values. This effect is most pronounced when the median reaches a local minimum in central tendency. To quantify this effect, we plotted the largest differences between the medians and the mean (referred to as undershoots) against the expected value λ (Fig 3A). The undershoots for the unbiased median (**+**) were fitted with an exponential function: *a*·exp(–λ/*τ*) – *b* (shown in green), where *a* = 0.0667±0.0019, *τ* = 1.41±0.11, *b* = 0.1507±0.0009 (here and further, the data are given as a mean ± SD). Similarly, the undershoots for the conventional median (**×**) were fitted with the same exponential form (shown in red), with parameters *a* = 0.0313±0.0012, *τ* = 1.50±0.16, *b* = 0.1608±0.0006. The average largest undershoot was smaller for the unbiased median compared to the conventional median significantly (–0.1381±0.0126 vs. –0.1544±0.0061, n = 5, *P* = 0.005, Student’s paired t-test). Furthermore, the difference between the baselines (–*b*) of the exponential fits was even more significant (−0.1507±0.0009 vs. −0.1608±0.0006, n = 5, *P* = 1.3·10^−8^, Student’s paired t-test). The key takeaway is that although both conventional and unbiased medians reduce the influence of extreme values, the unbiased median yields a more accurate estimate of the central tendency.

**Fig 3.**
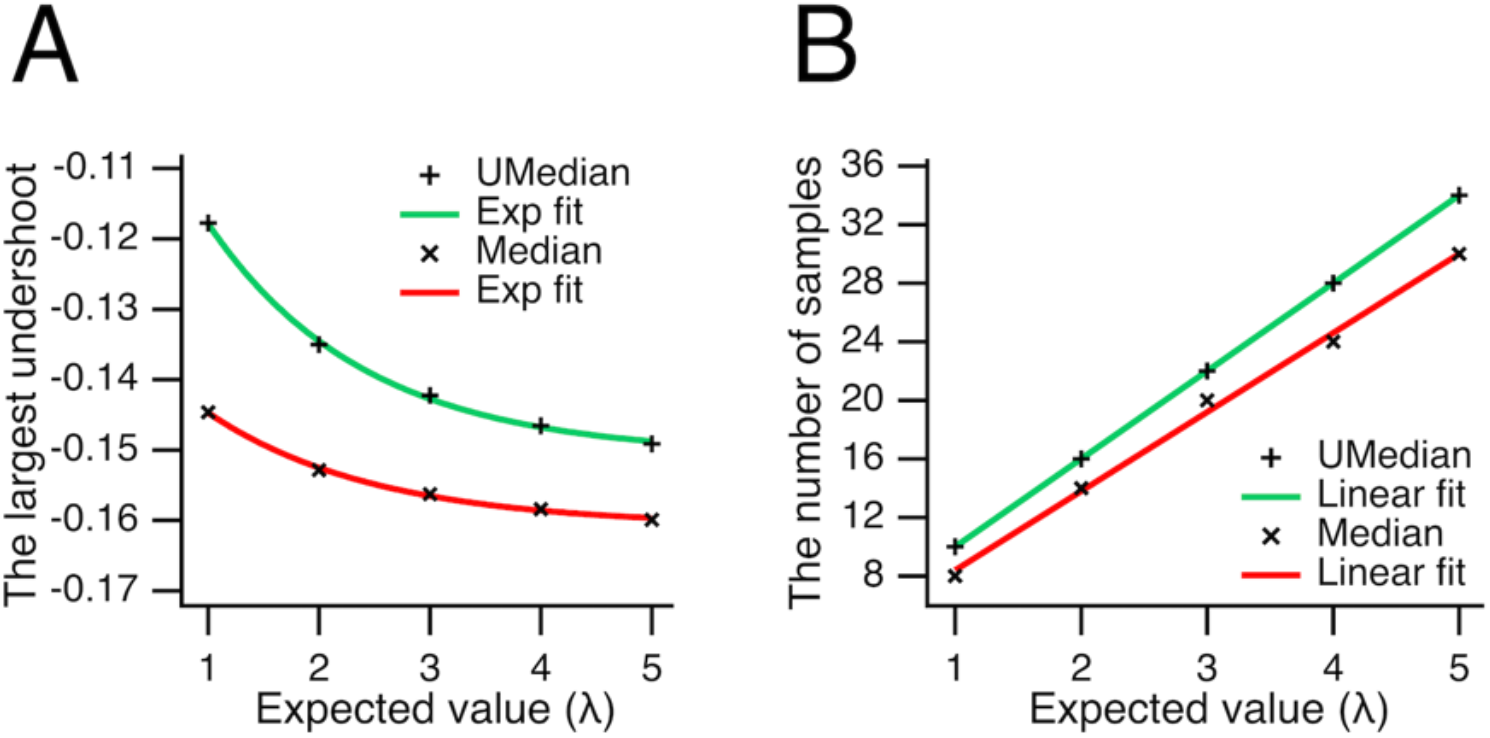
Reduced underestimation of the expected value by the unbiased median. (***A***) The largest difference (undershoot) between the estimated central tendency (using either the unbiased or conventional median) and the expected value (λ) is plotted against λ. Undershoots for the unbiased median (**+**) and conventional median (**×**) medians were fitted with the exponential *a*·exp(–λ/*τ*) – *b*. For the unbiased median (green), *a* = 0.0667±0.0019, *τ* = 1.41±0.11, *b* = 0.1507±0.0009. For the conventional median (red), *a* = 0.0313±0.0012, *τ* = 1.50±0.16, *b* = 0.1608±0.0006. On average, the largest undershoot was significantly smaller for the unbiased median than for the conventional median (–0.1381±0.0126 vs. –0.1544±0.0061, n = 5, *P* = 0.005, Student’s paired t-test). (***B***), The sample size at which the maximum undershoot occurs for the unbiased (**+**) and conventional (**×**) medians is plotted against λ. These values were fitted with a line *a*·λ + *b*. For the unbiased median (green): *a* = 6±0, *b* = 4±0. For the conventional median (red), *a* = 5.4±0.2, *b* = 3.00±0.66. Notably, the largest undershoot for the unbiased median consistently occurs at a smaller sample size than for the conventional median (*P* = 0.002, n = 5, Student’s paired t-test).

The sample sizes at which the largest undershoots occur in Poisson-distributed datasets are presented in Fig 3B. These sample sizes were fitted with linear functions of the form *a*·λ + *b*, yielding parameters *a* = 6±0, *b* = 4±0 for the unbiased median (shown in green), and *a* = 5.4±0.2, *b* = 3.00±0.66 for the conventional median (shown in red). A key finding is that the largest undershoot for the unbiased median consistently occurs at a smaller sample size compared to the conventional median (Student’s paired t-test, *P* = 0.002, n = 5).

To determine when the unbiased median offers the greatest advantage over the conventional median, we plotted the difference between these two estimators shown in Fig 2 against sample size (Fig 4A). The results indicate that this difference reaches its maximum at sample sizes of 4 or 6 (Fig 4A). Thus, the benefit of using the unbiased median—demonstrated previously in Fig 3—is particularly pronounced for small datasets containing 4 or 6 samples. This advantage becomes even more significant at lower expected values λ, as shown in Fig 4B: here, the largest combined difference between the medians at sample sizes 4 and 6 (♦) is plotted versus λ and fitted with the exponential function *a*·exp(–*λ*/*τ*) + *b*, where *a* = 0.0553±0.0007, *τ* = 0.409±0.024, and *b* = 0.02678±0.00003 (n = 2).

**Fig 4.**
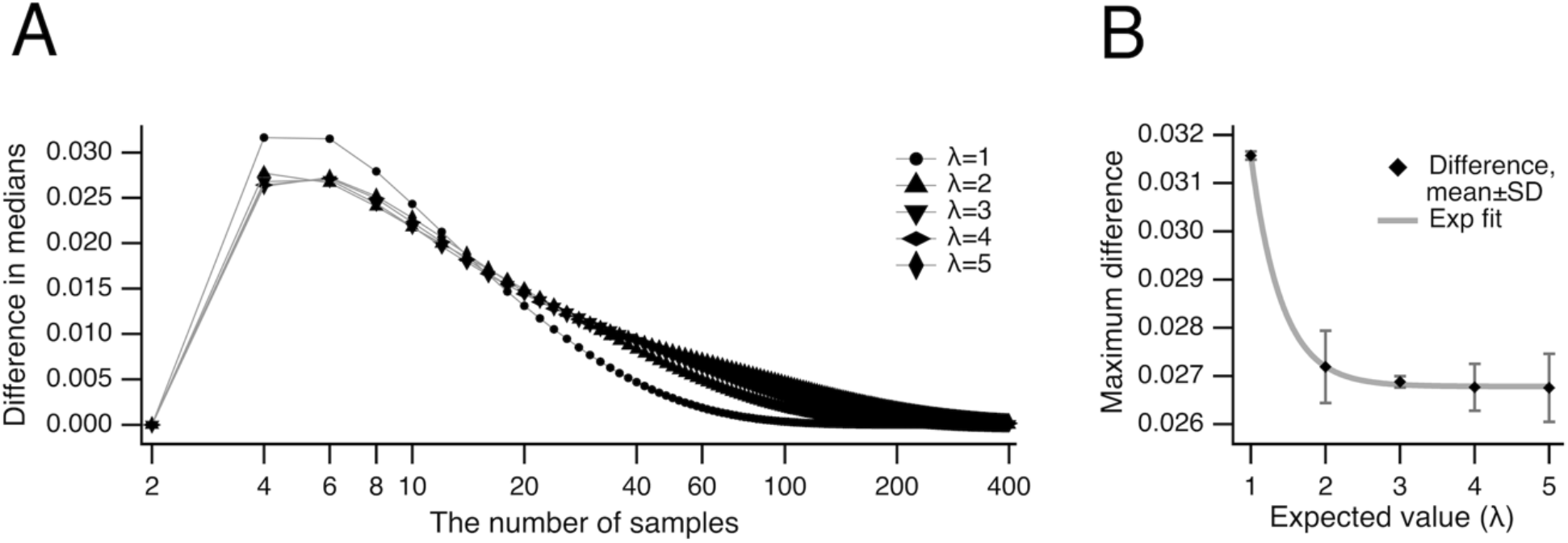
Peak difference between unbiased and conventional medians. (***A***) The differences between central tendency estimates using the unbiased and conventional medians are plotted for expected values λ ranging from 1 to 5. Note that these differences reach their maximum at sample sizes of 4 and 6. (***B***) The largest combined difference at sample sizes 4 and 6 (♦) is plotted against the expected value λ and fitted with an exponential function (grey): *a*·exp(–λ/*τ*) + *b*, where *a* = 0.0553±0.0007, *τ* = 0.409±0.024, and *b* = 0.02678±0.00003 (n = 2). Note that lower expected values correspond to greater differences between the two medians.

In data analysis, variance—which quantifies the spread or dispersion of values—is as essential a measure as central tendency, as it reveals how much data deviate from the average and enables assessment of reliability, consistency, and interpretability [10]. For Poisson-distributed datasets with sample size *n*, the theoretical variance is λ/*n*; thus, multiplying variance by *n* provides a convenient normalization for comparing results across different sample sizes. To compare the variances of the central tendency estimators—the unbiased median, conventional median, and mean—we normalized each variance by multiplying it by *n* and plotted these values against sample size (Fig 5). Normalized variances for central tendency measures in Poisson datasets, evaluated as the mean (black), conventional median (red), and unbiased median (green), are shown for expected values λ ranging from 1 to 5 (Fig 5). As expected for the Poisson process, the arithmetic mean (♦) closely matches the expected value λ (see Fig 2), with standard deviations from the expected value of 2.3·10^−4^, 5.3·10^−4^, 8.2·10^−4^, 1.1·10^−3^, and 1.3·10^−3^ for λ = 1, 2, 3, 4, and 5, respectively. These standard deviation values represent the lower bound of inaccuracy in subsequent analyses.

**Fig 5.**
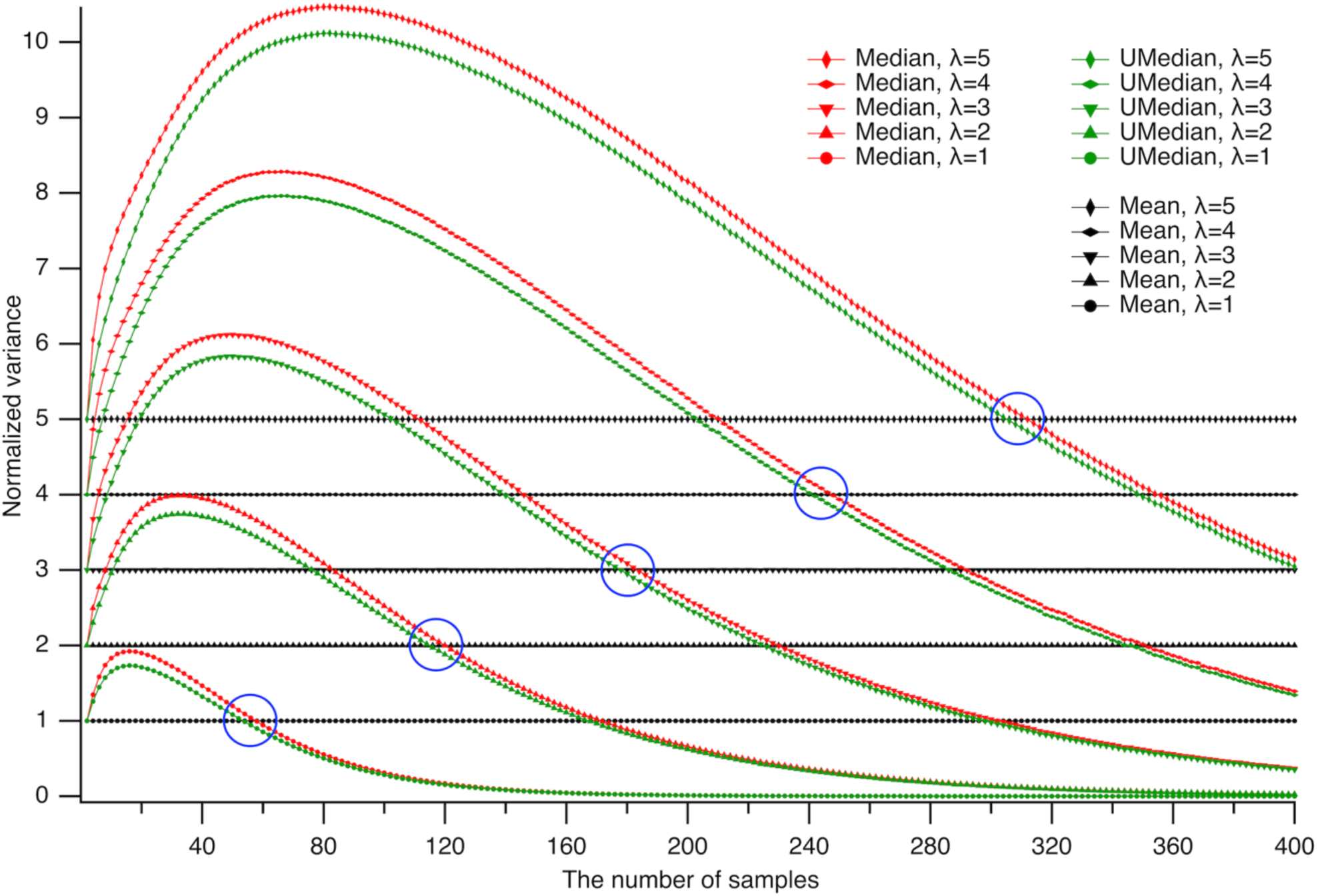
Variance overestimation by medians at small sample sizes. Normalized variances of central tendency estimators for Poisson-distributed datasets—computed as the mean (black), conventional median (red), and unbiased median (green)—are shown for expected values λ from 1 to 5. At small sample sizes, the variances of both median estimators exceed λ, with the unbiased median consistently exhibiting lower variance than the conventional median. As the sample size increases, the variances of both medians eventually drop below that of the mean, beyond sample-size thresholds indicated by blue circles—demonstrating the greater efficiency of median-based estimators at larger sample sizes. Notably, for the unbiased median, this crossover point occurs an average of 5.7 ± 1.1 samples earlier than for the conventional median (*P* = 0.0003, n = 5, Student’s paired t-test).

Note that at small sample sizes, the normalized variances of both medians overestimate the expected value λ, with the unbiased median consistently exhibiting lower variance than the conventional median. Notably, as sample size increases, the variances of both medians eventually fall below that of the mean beyond certain thresholds (as indicated with circles), highlighting the greater efficiency of median-based estimators for larger sample sizes. The earlier crossover point for the unbiased median (on average by 5.7 ± 1.1 samples, *P* = 0.0003, n = 5, Student’s paired t-test) demonstrates its greater efficiency compared to the conventional median.

Remarkably, at small sample sizes, the normalized variances of both median estimators exceed the expected value λ, with the unbiased median consistently displaying lower variance than the conventional median. As sample size increases, the variances of both medians eventually drop below that of the mean at specific thresholds (indicated by circles in Fig 5), illustrating the greater efficiency of median-based estimators for larger datasets. Importantly, the unbiased median achieves this crossover point significantly earlier—by an average of 5.7 ± 1.1 samples (*P* = 0.0003, n = 5, Student’s paired t-test)—confirming its superior efficiency compared to the conventional median.

It may seem counterintuitive that the variances of the medians overestimated the variance of the mean, since one might expect that the medians’ tendency to disregard extreme values in Poisson datasets would reduce, rather than increase, their variance. However, when recognizing that, at small sample sizes, the medians systematically underestimate central tendency (Fig 2), the corresponding overestimation of the deviation from the central tendency (Fig 5) is a logical consequence rather than an unexpected result. The advantage of the medians in disregarding extreme values becomes evident at large sample sizes: the central tendency estimates converge to the expected value, and the variance of the medians falls below that of the means beyond the encircled regions in Fig 5.

Normalizing the variances in Fig 5 facilitated comparison across large sample sizes. However, for practical data analysis, it is essential to also consider the absolute (non-normalized) variances, as their magnitudes provide direct information about variability in the original measurement scale. To quantify how much the variance estimates from the medians exceed that of the mean, we plotted the difference between the variances of the medians and the variance of the mean as a function of sample size (Fig 6A). The relative variance estimate from the conventional median (shown in red) peaked consistently at a sample size of 6 across all expected values λ. In contrast, the unbiased median (shown in green) exhibited peak relative variance at sample size 6 for λ = 1 and 5; at sample size 8 for λ = 3 and 4; and at sample size 10 for λ = 2. The peaks in the relative variances represent the points of greatest inefficiency of the medians compared to the mean. To illustrate this, we plotted the peak values (referred to as overshoots) against the expected value λ (Fig 6B). These overshoots were fitted with a quadratic polynomial, *a*·λ^2^ + *b*·λ + *c*, yielding parameters: for the conventional median (red), *a* = 0.0024±0.0009, *b* = 0.0297±0.0055, *c* = 0.0636±0.0073, and for the unbiased median (green), *a* = 0.0008±0.0002, *b* = 0.0177±0.0013, *c* = 0.0561±0.0017. On average, the largest overshoot was significantly smaller for the unbiased median compared to the conventional median (0.1186±0.0361 vs. 0.1788±0.0696, n = 5, *P* = 0.016, Student’s paired t-test), demonstrating the superior performance of the unbiased median.

**Fig 6.**
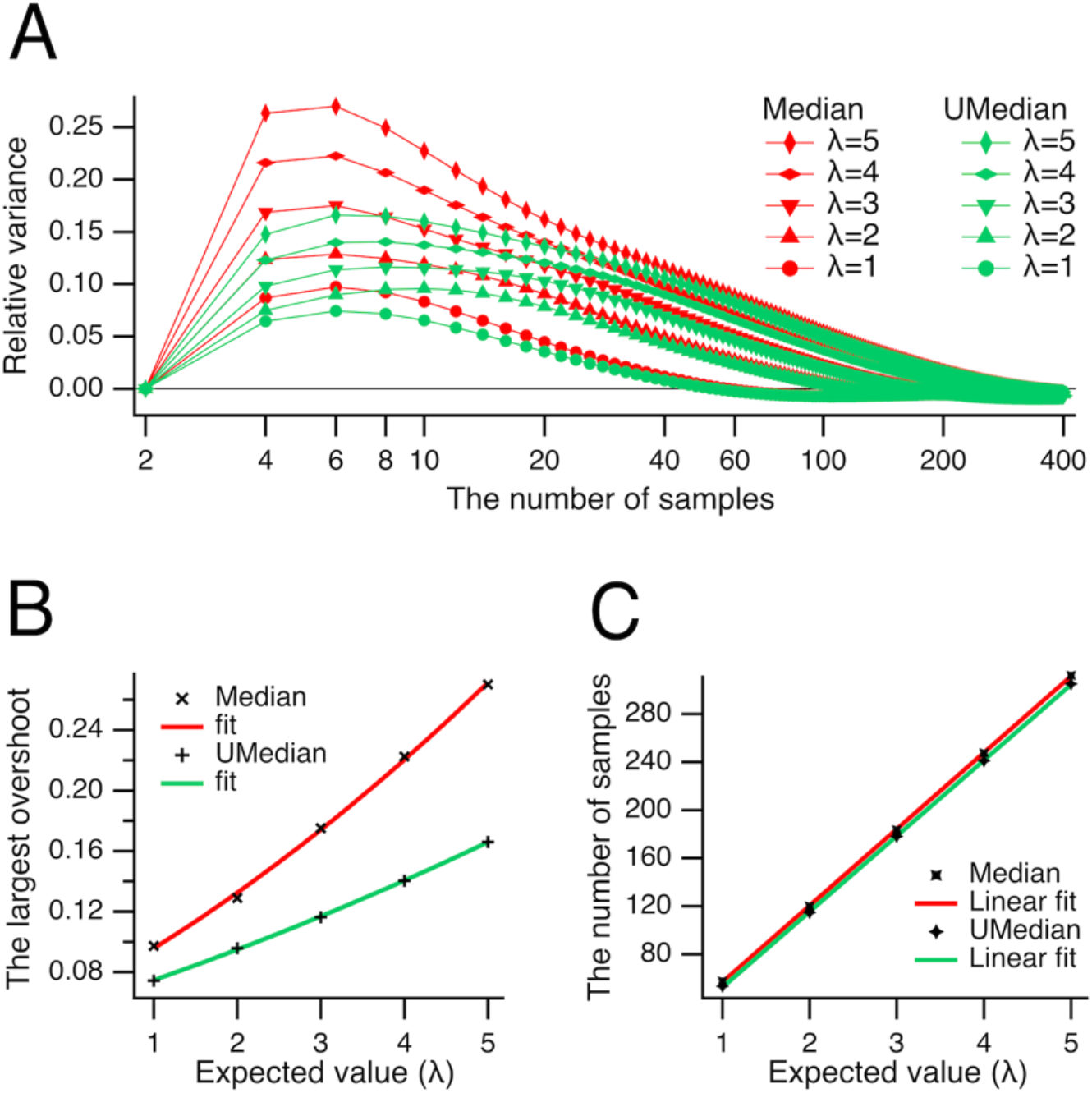
Peaks and zeros of the relative variances. (***A***) The relative variance estimate from the conventional median (shown in red) peaked at the sample size of 6 at all expected values λ, whereas that of the unbiased median (shown in green) peaked at samples sizes of 6 (at λ = 1, 5), 8 (at λ = 3, 4), and 10 (at λ = 2). (***B***) The largest overshoots for the conventional (**×**) and unbiased (**+**) medians are plotted against the expected value λ. The overshoots were fitted with a quadratic polynomial *a*·λ^2^ + *b*·λ + *c*, yielding parameters: for the conventional median (red), *a* = 0.0024±0.0009, *b* = 0.0297±0.0055, *c* = 0.0636±0.0073, and for the unbiased median (green), *a* = 0.0008±0.0002, *b* = 0.0177±0.0013, *c* = 0.0561±0.0017. On average, the largest overshoot was significantly smaller for the unbiased median than for the conventional median (0.1186±0.0361 vs. 0.1788±0.0696, n = 5, *P* = 0.016, Student’s paired t-test). ***C***, The number of samples at zero relative variances of conventional (**×**) or unbiased (**+**) medians. The sample sizes were fitted with a line *a*·λ – *b*, where *a* = 63.63±0.26, *b* = 6.78±0.85 for conventional median (red), and *a* = 62.95±0.27, *b* = 10.43±0.90 for unbiased median (green).

Notably, the relative variances were highest for sample sizes between 8 and 34, coinciding with the greatest underestimation of central tendency by the medians (see Figs 2 and 3C). As the medians approached the mean at larger sample sizes (Fig 2), the corresponding relative variances also dropped below zero at the sample sizes indicated in Fig 6C. The sample sizes were fitted with a line *a*·λ – *b*: for the conventional median (red), *a* = 63.63±0.26, *b* = 6.78±0.85; for the unbiased median (green), *a* = 62.95±0.27, *b* = 10.43±0.90.

One can observe that the relative variance estimate of the unbiased median was consistently lower than that of the conventional median for the same expected value λ (Figs 6A,B), indicating a clear advantage in using the unbiased median. To identify the conditions under which the unbiased median provides the greatest benefit, we analyzed the difference between the variances of the unbiased and conventional medians (Fig 7A). This difference was the largest at sample sizes of 4 and 6. To further explore how this difference varies with sample size, we plotted the combined variance difference at sample sizes 4 and 6 against the expected value λ (Fig 7B). The largest difference for the number of samples 4 and 6 (combined, n = 2) was fitted with a line *a*·λ – *b*, where *a* = 0.0219±0.0001, *b* = 0.0004±0.0003.

**Fig 7.**
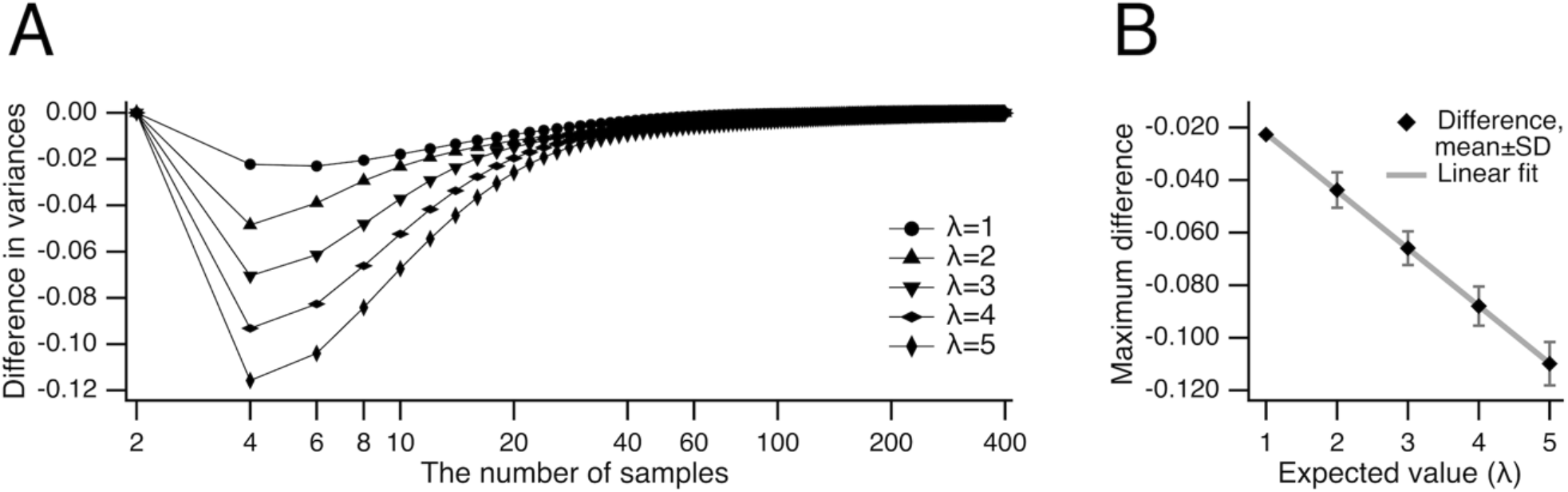
Difference in variances of unbiased and conventional median. (***A***) Difference between the variances of central tendency of Poisson’s datasets evaluated as unbiased and conventional medians are shown for expected values λ from 1 to 5. Note that the difference is the largest at the number of samples 4 and 6. (***B***) The largest difference (♦) shown in panel (***A***) for the number of samples 4 and 6 (combined, n = 2) for expected values λ from 1 to 5 is fitted with a line *a*·λ – *b*, where *a* = 0.0219±0.0001, *b* = 0.0004±0.0003.

## Discussion

In this study, we proposed a novel median estimator for even samples sizes that, unlike the conventional median, is unbiased for asymmetrical distributions. This unbiased estimator was developed by re-framing the original *ill-posed* problem as a special case of a *well-posed* one: “finding the median by minimizing the sum of deviations raised to a power that approaches unity”. Compared to the conventional median, the proposed median is a more accurate, lower-variance estimator of central tendency for asymmetrically distributed datasets.

We refer to the proposed median estimator as the *unbiased* median to distinguish it from the conventional median. The term “unbiased” was chosen for two main reasons. First, the conventional median—defined as the arithmetic mean of the two middle values—implicitly assumes that the data distribution is symmetrical, thereby introducing bias when this assumption does not hold. In contrast, the unbiased median, as calculated from equation (11), makes no such assumption and is therefore free from bias toward symmetrical distributions. Second, the name is an analogy to the concept of bias in statistics. For instance, the sample variance formula uses a correction (dividing by *n* − 1) because using the sample mean would otherwise create a biased estimate of the true population variance. In a similar way, the conventional median can be seen as a biased estimator of the population mean if the data’s distribution is asymmetrical.

In the Poisson distribution (equation 17), *k* represents the number of events per measurement interval and must therefore be an integer. However, the expected number of events λ—defined as the arithmetic mean—can take any positive rational value. We assessed the properties of medians for Poisson-distributed datasets at discrete expectation values (λ = 1, 2, 3, 4, and 5). In practice, however, λ can range from values near zero to very large numbers, depending on the underlying process. To enable readers to interpolate between these discrete values and, where appropriate, extrapolate beyond them, we fitted the data with suitable functions (see Figs 3, 4, 6, and 8). For example, extrapolating the exponential fits from Fig 3A predicts that the maximal difference between the central tendencies measured by the medians occurs at λ ≪ 1 and a sample size of four (see Fig 4A).

**Fig 8.**
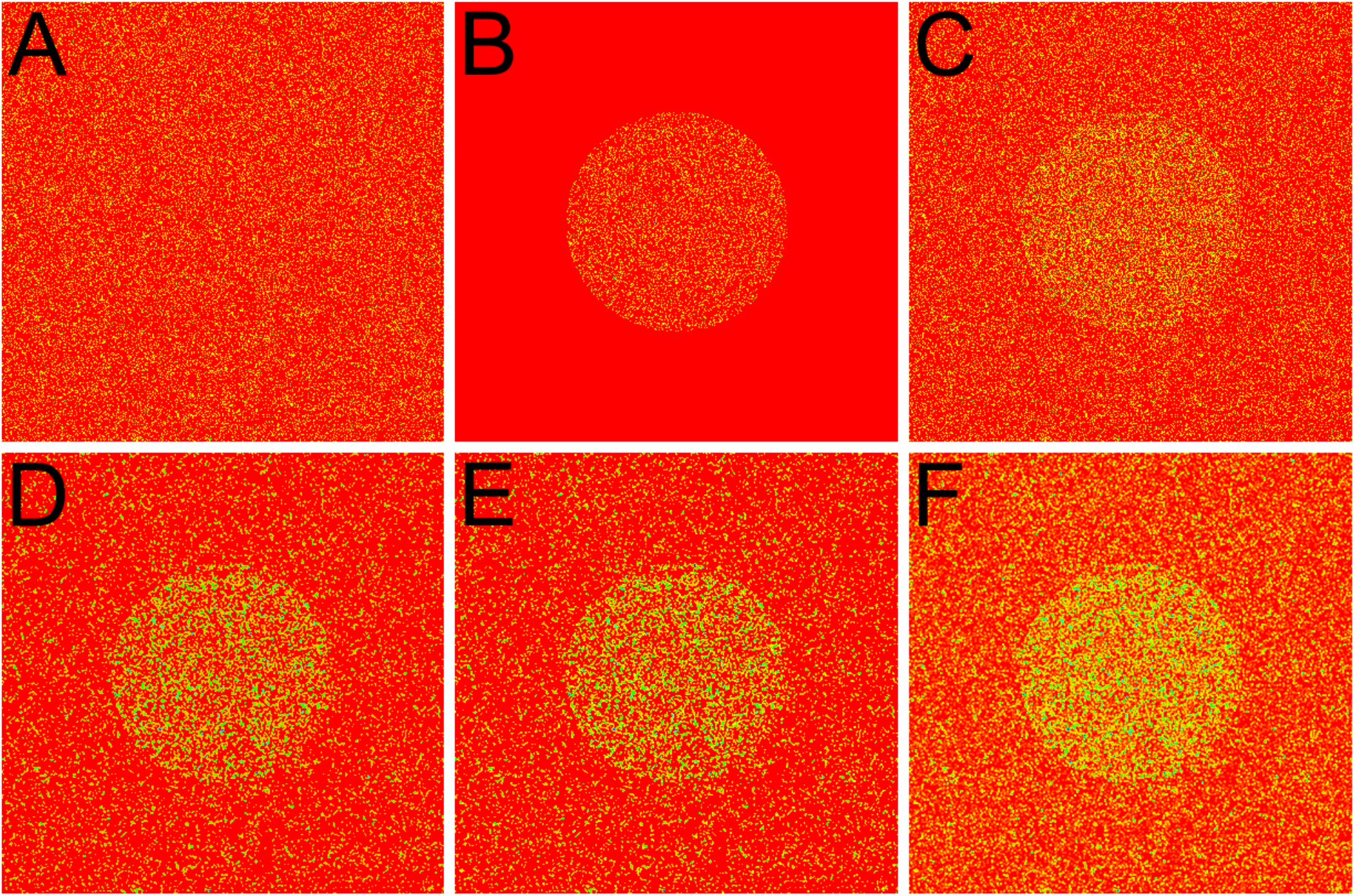
Qualitative advantage of the unbiased median. For clarity, grayscale images were mapped to a rainbow color scale: red denotes the lowest gray values, yellow to green correspond to intermediate values, and blue indicates the highest values. (***A***) Dark current image of the photodetector array (expected value λ = 1/5). (***B***) Incident photon flux shaped as a circle (λ = 1/5). (***C***) Combined output of the photodetector array (sum of dark current and photon flux). Panels ***A, B***, and ***C*** are all scaled to the same maximum value (the highest pixel value in panel ***C***). Panels ***D, E, F*** show the results of filtering the image from panel ***C*** using a 2×2 neighborhood with the conventional median (***D***), unbiased median (***E***), and mean (***F***), respectively. Images in ***D, E***, and ***F*** are scaled to the maximum value in panel ***F***. Note that brightness and contrast increase from the conventional median (panel ***D***) to the unbiased median (panel ***E***), and to the mean (panel ***F***).

While the difference between the central tendencies measured by the unbiased and conventional medians is statistically significant, it remains to be determined whether this difference is substantial enough to be perceptually meaningful. In other words, does a quantitative difference result in a qualitative change? To address this, we simulated the capture of an image of a circle using a 2D photodetector array. Since the maximal difference between the medians occurs at λ ≪ 1 and a sample size of four (see Figs 3A and 4A), we set the expected values for both photon flux and photodetector dark current to λ = 1/5. The resulting images were then filtered using the conventional median, the unbiased median, and the mean of 2×2 neighboring pixels. For enhanced visualization, the grayscale images were mapped onto a rainbow color scale: red indicates the lowest gray values, yellow to green corresponds to intermediate values, and blue represents the highest values. Fig 8A displays the dark current of the photodetector array; Fig 8B shows the incident photon flux; and Fig 8C presents the combined image (sum of dark current and photon flux). Panels ***D, E, F*** represent the image from panel ***C*** being filtered using conventional median, unbiased median, and mean, respectively. The images in panels ***A, B***, and ***C*** are all scaled to the same maximum value (the highest pixel in panel ***C***); similarly, images in panels ***D, E***, and ***F*** are scaled to the maximum value in panel ***F***.

Remarkably, all filtered images in panels ***D, E***, and ***F*** reveal the circle with much greater clarity than the unfiltered image in panel ***C***. Brightness and contrast progressively increase from the conventional median (panel ***D***), to the unbiased median (panel ***E***), and finally to the mean (panel ***F***). As expected, the mean produces the brightest image, since both median estimators tend to underestimate the mean (Figs 2 and 3). However, the unbiased median produces a distinctly brighter image than the conventional median, underscoring its reduced underestimation of the mean. These results demonstrate that the unbiased median provides a more accurate estimate of central tendency than the conventional median. Thus, the quantitative improvement is sufficient to result in a qualitative difference that is perceptible to the naked eye.

Strictly speaking, using medians to analyze Poisson-distributed data is not ideal, since such data inherently lack outliers [11]. By disregarding extreme positive values—which are a natural part of the Poisson distribution—medians tend to underestimate the expected value (see Figs 2 and 3) and, as a consequence, overestimate the variance (see Figs 5 and 6; also: [12, 13]). In real-world datasets, it is often difficult to determine whether outliers are present. For instance, in scenarios where outliers are unlikely—such as measuring photon counts from a photodetector with zero dark current—the median systematically underestimates the central tendency, as illustrated in Figs 2 and 3. Conversely, when genuine outliers are present and serve as the primary source of asymmetry in the data, using the median to estimate central tendency becomes particularly advantageous—especially under the similar conditions in which it exhibits the greatest underestimation for Poisson-distributed data.

The only limitation of the unbiased median is that, for sample sizes of ten or more, its calculation requires iterative methods based on equation (11), making it considerably more computationally demanding than the conventional median, which is obtained simply by averaging the two middle values.

## Conclusion

Overall, the results demonstrate the clear advantage of using the unbiased median over the conventional one: (i) the unbiased median (equation 11) provides a unique solution to a well-posed problem, free from any bias toward symmetrical distributions; (ii) central tendency of asymmetrically distributed data measured as unbiased median is more accurate (Figs 2, 3A, 4A); (iii) the variance of central tendency measured as unbiased median is lower (Figs 5, 6A, 7A); (iv) the superior results provided by the unbiased median can be evident upon visual inspection (Fig 8).

## Acknowledgements

This study was funded by grant R01AG066905 to S.A. Thomas from the National Institutes of Health.

## Notes

### Competing Interest Statement

The authors have declared no competing interest.

### Summary of Updates

Figure 7 Legend has been updated.

